# stainlib: a python library for augmentation and normalization of histopathology H&E images

**DOI:** 10.1101/2022.05.17.492245

**Authors:** Sebastian Otálora, Niccoló Marini, Damian Podareanu, Ruben Hekster, David Tellez, Jeroen Van Der Laak, Henning Müller, Manfredo Atzori

## Abstract

Computational pathology is a domain of increasing scientific and social interest. The automatic analysis of histopathology images stained with Hematoxylin and Eosin (H&E) can help clinicians diagnose and quantify diseases. Computer vision methods based on deep learning can perform on par or better than pathologists in specific tasks [1, 2, 15]. Nevertheless, the visual heterogeneity in histopathology images due to batch effects, differences in preparation in different pathology laboratories, and the scanner can produce tissue appearance changes in the digitized whole-slide images. Such changes impede the application of the trained models in clinical scenarios where there is high variability in the images. We introduce `stainlib`, an easy-to-use and expandable python3 library that collects and unifies state-of-the-art methods for color augmentation and normalization of histopathology H&E images. `stainlib` also contains recent deep learning-based approaches that perform a robust stain-invariant training of CNN models. `stainlib` can help researchers build models robust to color domain shift by augmenting and harmonizing the training data, allowing the deployment of better models in the digital pathology practice.

## 1. Motivation and significance

During the last decade, the automatic analysis of digital pathology images has increased until the point where commercial products are now available, and new ones are being cleared by health control and supervision organizations. In research, modern computational pathology techniques are based on the steady development of deep learning algorithms and deep convolutional neural networks (CNN) [13, 9]. Shifting from handcrafted features towards end–to–end training of deep learning models made it possible to automatically detect cancer in digitized histopathology images, both, in image regions and at the whole-slide-image level, with performances previously unseen [1, 2].

Methods have become more reliable, achieving in some cases a performance that is comparable to pathologists for specific segmentation and classification tasks [1, 5, 2, 15, 9, 12]. Despite the remarkable good performance of some methods, there are still technical barriers that prevent the translation of these advances into clinical applications [17]. A clinically applicable deep learning method needs to be able to cope with the heterogeneity in the color of the images that arises from preparing and staining the tissue samples in the pathology laboratory [14, 19]. Stains are chemical reagents that attach to specific proteins and that are used to enhance the contrast between different tissue structures for their examination under a microscope by a pathologist. The most commonly used stains are a combination of hematoxylin and eosin (H&E). These two reagents highlight the nuclei DNA content with a dark blue-purple color (Hematoxylin) and cytoplasm and stromal matrix contents with a light pink-red color (Eosin), see Figure 1 for exemplar H&E stained image regions of prostate tissue.

**Figure 1:**
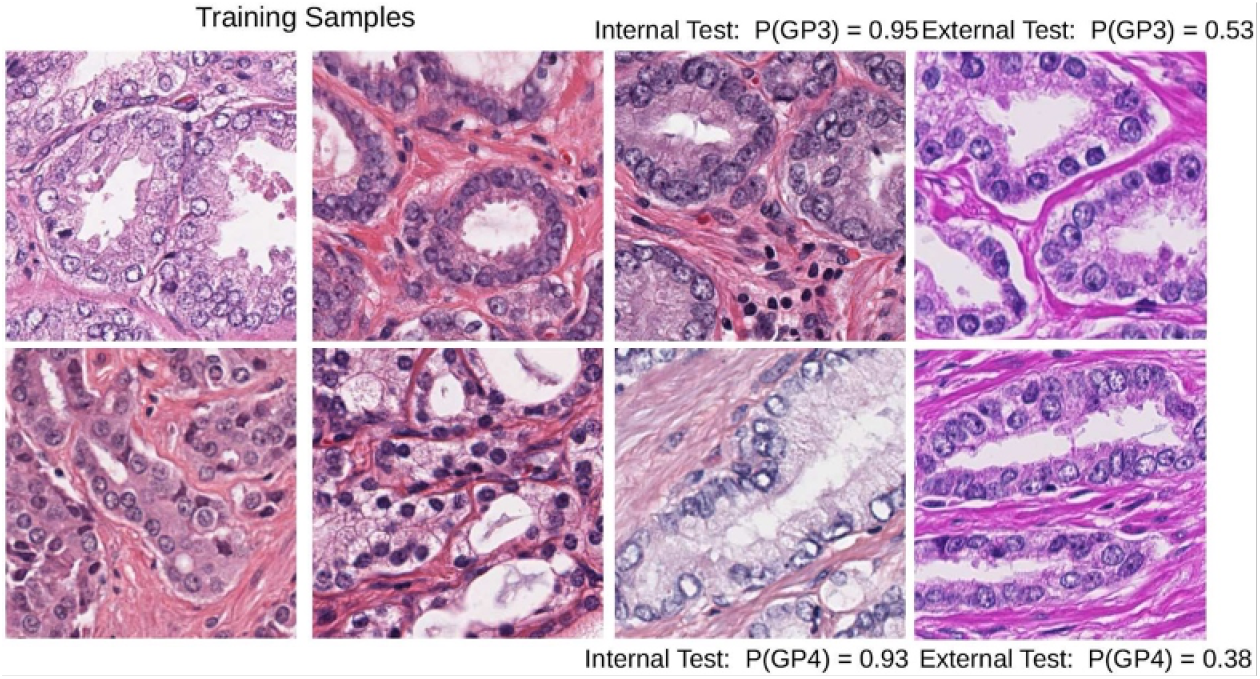
Training images with H&E concentrations that are noticeable different from external test sets can lead to model misclassifications; image taken from [14]. P(GP3) and P(GP4) stands for probability of the image region to contain Gleason patterns 3 and 4 respectively.

One of the most important factors preventing the application of machine learning methods in clinical practice is related to the heterogeneity of H&E images due to the many parameters involved in the tissue preparation and the digital scanning process (temperature of the tissue, the thickness of the cuts, the image sensor of the digital camera, the stitching algorithm among others) [11]. Figure 1 shows an example of stain variation in training and test sets and its impact on the performance of a CNN model trained only with partial variations. Two approaches are most commonly applied to take into account such variations when training CNN models. First, methods that transform an input H&E image given a target image template are known as “stain normalization”. Their aim is to match the input color distributions (or H&E concentrations) with the one given in a target image. The second approach refers to “stain” or “color augmentation methods”, which create new synthetic samples to increase the training dataset size, creating more robust models regarding color variations. There are novel image processing and machine learning techniques reported in the literature to deal with color heterogeneity, improving classification, and segmentation performance for various tissue types [19, 6, 14, 4, 11]. While the specific normalization technique depends on the task to solve [19, 4], recent work has reported consistent improvements in performance and robustness to external datasets employing color augmentation techniques [19, 4] or a combination of normalization and augmentation [14].

There are existing tools and methods to deal with H&E color heterogeneity, however not unified under a standard tool-set, many are also written in different programming languages, with various dependencies, and others are unknown by some researchers.

Few libraries comprise multiple methods for H&E image normalization and augmentation. Furthermore, only a handful of methods tackle color heterogeneity in H&E images using the modern machine and deep learning techniques. The codebase from articles in the literature is mostly in self-contained repositories, and its evaluation is usually performed in ad-hoc tasks using specific and often private datasets. The lack of libraries with multiple methods limits the possibilities to evaluate the best strategy to deal with color heterogeneity for new datasets or tasks.

This article aims to present and validate stainlib, an easy-to-use, extensible library to extract homogeneous representations of heterogeneous color information. With stainlib we make an effort to find, extract, collect, test, and unify most of the existing methods into a single library and make them easy to use. stainlib includes the most commonly used methods for color augmentation and normalization of histopathology images, having input local image regions (or patches). It contains classical machine learning and novel deep learning techniques to tackle the heterogeneity of color in H&E images.

### 1.1. Related work

QuPath, Staintools, and HistomicksTK, are likely among the most popular existing software tools to deal with color heterogeneity. QuPath (https://qupath.github.io/.) is an open and extensible software platform for Whole Slide Image (WSI) analysis. It includes methods for estimating and setting stain vectors. Scripts created for running specific color normalization methods can also be used within QuPath. Due to the big codebase of QuPath, it is challenging to run a classification or segmentation model without having to write a considerable amount of scripts to have a full pipeline, taking into account datasets with considerable color heterogeneity.

Staintools is a set of tools for tissue stain normalization and augmentation in python 3. It contains implementations that follow the same coding style of scikit-learn, where the methods are made to fit or train a model. It is open-source and can be downloaded from the Github repository: https://github.com/Peter554/StainTools. The library contains two extractive normalization methods (Macenko, Vahadane), and the only augmentation techniques included in staintools are based on the same extractive methods by modifying the estimated stain concentrations.

HistomicksTK is a python toolkit for histopathology image analysis. It contains several methods for stain normalization and color augmentation based on the stain perturbation methods from Tellez et al. [18]. It can be downloaded from https://pypi.org/project/histomicstk/. The HistomicksTK toolkit contains many overlapping methods with stainlib but it lacks modularity to use the color tools as standalone modules, which creates difficulties for its usage in different research scenarios. Despite the existence of few libraries, the domain still lacks a modular library including both standard and more modern methods that can be easily evaluated on varying datasets.

## 2. Software description

Stainlib is a python library containing methods for H&E image normalization and augmentation. The objective is to develop an easy-to-use python3 library that includes the most commonly used methods for color augmentation and normalization of histopathology images, having as input local image regions and to add more recent methods to tackle color heterogeneity based on deep learning approaches, too. In Figure 2, the structure and methods included in the library are displayed. The library can be downloaded from the following github repository: https://github.com/sebastianffx/stainlib.

**Figure 2:**
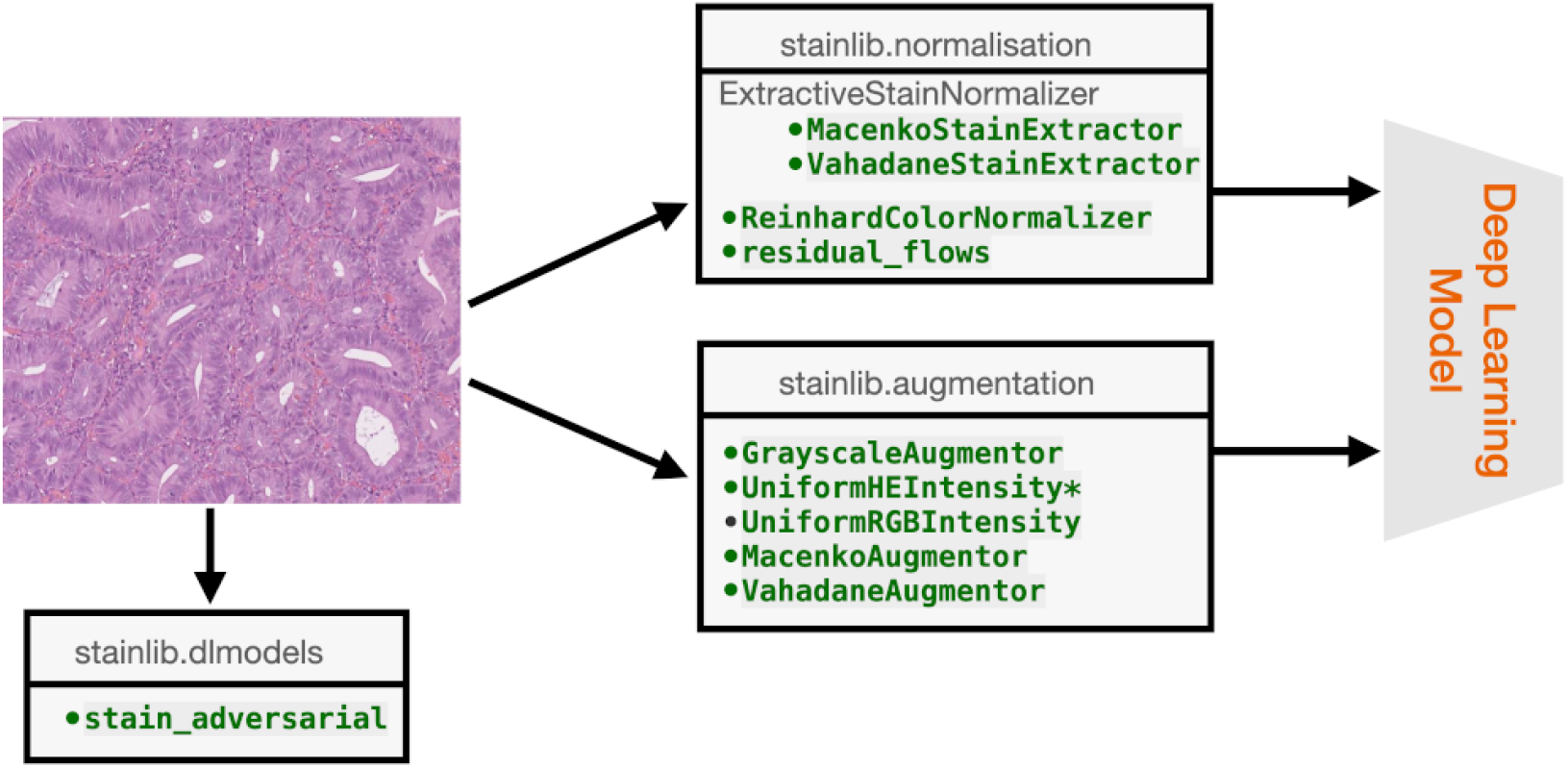
Implemented methods in stainlib for stain normalization and augmentation. The first version of the library includes all the methods in green and linked submodules of the residual flows and stain adversarial methods. For next versions of the library, further state of the art methods will be included.

### 2.1. Software Architecture

`stainlib` is composed of three main modules `stainlib.augmentation`, `stainlib.normalization`, `stainlib.dlmodels`. The methods and the underlying theory of each of the modules are explained in the following subsections.

#### 2.1.1. stainlib.normalization

In digital pathology, the thin slice tissue cuts that are counter-stained in the H&E tissue slides are digitized using digital tissue scanners. The stained tissue slide’s light absorbance is quantified and represented in a computer as a two-dimensional digital image (despite coming from a three-dimensional biological structure). In general, if the digital representation of the image is in the RGB color space, each pixel should contain a composition of the color representation of Hematoxylin (*purple*), Eosin(*pink*), and background (*white*).

Images acquired from the same center and using the same preparation methods share similar stain absorbance coefficients, which can be written as the linear transformation (omitting background that should be close to white for the three channels):

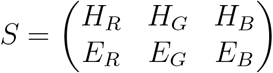

Where the first-row vector corresponds to the RGB components of hematoxylin and the second one to the components of Eosin. In staining normalization methods, the aim is to estimate the individual staining absorbance coefficients of the images *S* and quantify the absorbed light **C** by the tissue when it is scanned, which is the value in the H&E space of each pixel. The Beer-Lambert law provides a way to estimate them in the optical density space, given the original pixel content for the *c*-channel *I_c_*:

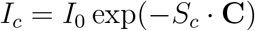

Where *c* ranges in the RGB channels, *S* ∈[0, +∞]^3×2^is the matrix of absorbance coefficients, **C** ∈ [0, +∞]^2^is the vector of the two staining concentration coefficients and *I*_0_ is the background value.

Several well-known stain extraction methods provide an estimation of *S*. In the widely used method of Macenko [10], this matrix is computed by calculating a plane using the two largest singular value decomposition vectors of the image and then projecting the data into this plane and clipping extreme values. In the method of Vahadane [20] this estimation is done by learning a sparse non-negative matrix factorization.

An alternative approach is to use residual flows for invertible generative modeling [3]. Flow-based generative models parameterize probability distributions through an invertible transformation and can be trained by maximum likelihood. Invertible residual networks provide a flexible family of transformations where only Lipschitz conditions rather than strict architectural constraints are needed for enforcing invertibility.

In `stainlib` we include the Vahadane and Macenko methods using the code base from the implementations in the staintools library^1^. In the Reinhard method [16], the color histogram of the source image (in the LAB color space) is matched with the target image. Despite its simplicity and original domain of application of natural images, it yields good results in histopathology images. We have also included the Reinhard normalization method in `stainlib`.

The invertible flows method is fully compatible with `stainlib` and can be used from the base implementation ^2^.

#### 2.1.2. stainlib.augmentation

It is now well known that deep learning classification and segmentation models for histopathology yield better results when data augmentation is used [19, 6, 14, 7]. The benefits of data augmentation might be intuitive in training deep learning models, where the larger the amount of data the model is fed with, the more variations the model is exposed to. Therefore, data augmentation usually can make the model more robust to changes in appearance in the test set. When there is a wide range of images with variations in color and preparation sources included in the training set, the models are more likely to output the correct prediction for new samples. Such a range of variations could be synthetically simulated, especially for the color variations in tissue appearance due to the stain concentrations. In color-information `stainlib` we have included five stain augmentation methods: 1) grayscale transformation, 2) shifts of the stain concentrations in H&E space, 3) shifts of the stain concentrations in RGB space, 4) shifts of the stain concentration matrix using the Macenko method and 5) shifts of the stain concentration matrix using the Vahadane method.

For the RGB to grayscale transformation, the method `rgb2gray` from the `scikit-image` library is used to generate realistic samples using random uniform variations of the grayscale values within a fixed range as follows:

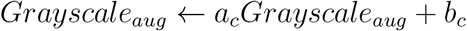

Where *a_c_* and *b_c_* are values drawn from a uniform distribution in the [0, 1] range. Similarly, for the shifts of the RGB channels, we generate new samples as follows for each channel, *c* as follows:

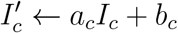

Again, *a_c_* and *b_c_* are values drawn from a uniform distribution in the [0, 1] range. In the case of the Macenko and Vahadane augmentation, we use the estimated stain H&E concentration matrices of the methods and shift them as follows:

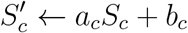

Where the values *a_c_* and *b_c_* are drawn from a uniform distribution in the [1 – *α_c_*, 1 + *α_c_*] and [1 – *β_c_*, 1 + *β_c_*] range. Values of *α* = 0.2 and *β_c_*= 0.2 were set as default values as they usually yield good qualitative and quantitative results in experiments. In the case of the augmentations based on the shifts of the stain concentrations in H&E space, we followed the implementation of Tellez et al. [18]. Section 3 presents a qualitative evaluation of the normalization and augmentation methods included in `stainlib`. Quantitative evaluation of these methods has also been performed in previous research from the authors [14, 6, 19].

#### 2.1.3. stainlib.dlmodels

Domain-invariant training of CNN’s is a promising technique to address training a single model for different domains. It includes the source domain information to guide the training towards domain-invariant features, allowing to achieve state-of-the-art results in classification tasks. In the case of training classification models with histopathology images, the domain represents the center where the tissue preparation characteristics are similar, e.g., hospital A, hospitals B. This technique shows excellent generalization performance to external test sets, and further improvements have been reported, when combined with data augmentation techniques [14, 8].

To explicitly write all the possible variations that lead to changes in the appearance of H&E images is infeasible. Stain invariant training of CNN models aims at detaching the domain or center information, where the changes in appearance originate, from the features that the model learns. In Figure 3, the inner workings of a stain invariant model is demonstrated by showing the flow of gradients in a small neural network example. The CNN has shared features for both the mitosis/no-mitosis classifier and the domain classifier. The main difference with a multi-task CNN is that the gradients from the domain branch are reversed to allow the penalization of unwanted domain information in the features. The stain-invariant model penalizes when the learned features help classify the domain, guiding the model towards features that do not consider the domain information. Evaluation of stain invariant models is described in detail in a previously published journal article from the authors [14].

**Figure 3:**
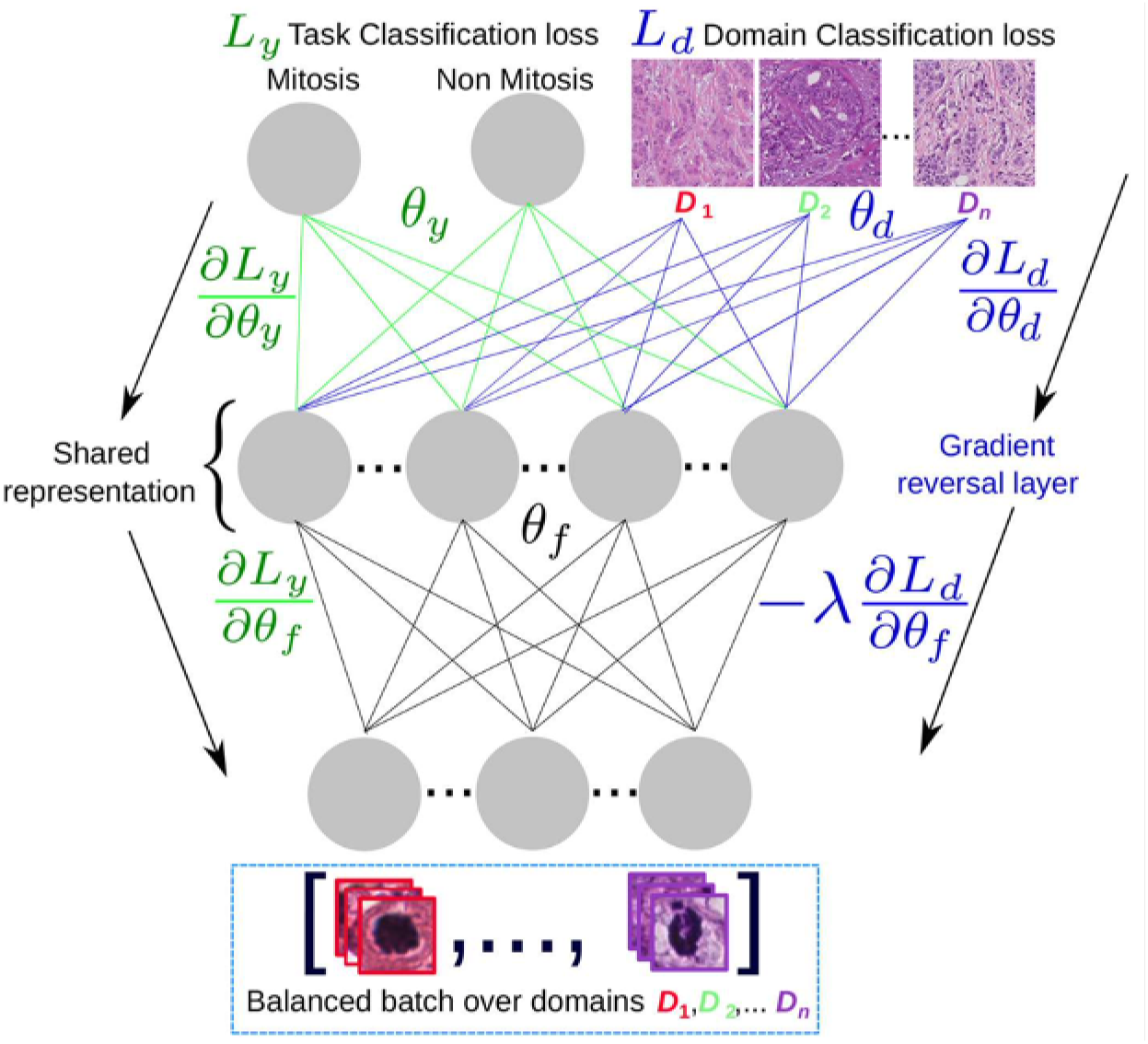
Domain adversarial scheme: A domain-balanced batch of images is passed as input to the network that has two types of outputs: the task classification output and the domain classification output. The shared representation *θ_f_* is optimal for the task classification and unable to discriminate between the *n* domains.

### 2.2. Software Functionalities

- **H&E image data augmentation**: This functionality allows generating one or more synthetic, yet realistic, copies of an image region extracted from a H&E WSI. Currently, five augmentation techniques are implemented, as described in section 2.1.2.
- **H&E image normalization**: The second main functionality is to normalize the color of image regions extracted from a H&E WSIs, given a template image. Currently, four normalization techniques are implemented, as described in section 2.1.1.
- **Stain invariant training of CNNs**: This functionality refers to the enabling of training a deep convolutional neural network with H&E images to tackle the stain heterogeneity when the center or source of the images is known. This source-code includes the python implementations of gradient reversal strategies [8, 14] and examples in a CNN model for the classification of H&E images of prostate and breast tissues. The method is explained and illustrated in Section 2.1.3.

## 3. Visual Examples

To demonstrate the proposed capabilities of the library, we show examples for augmentation and normalization using openly accessible images. The code snippets necessary to reproduce these examples are contained in a jupyter notebook in the source-code repository.

### 3.0.1. Image Augmentation

In Figure 4 the first row corresponds to ten grayscale augmented versions of the leftmost image using the method described in section 2.1.2. The second row shows ten augmented images using the light-HED augmentation from Tellez et al. [18]. Finally, the third and fourth rows correspond to the augmentation methods based on Macenko and Vahadane techniques, respectively. The augmented versions are generated perturbing the estimated stains, as in the case for stain normalization. Here the images for both methods look similar.

**Figure 4:**
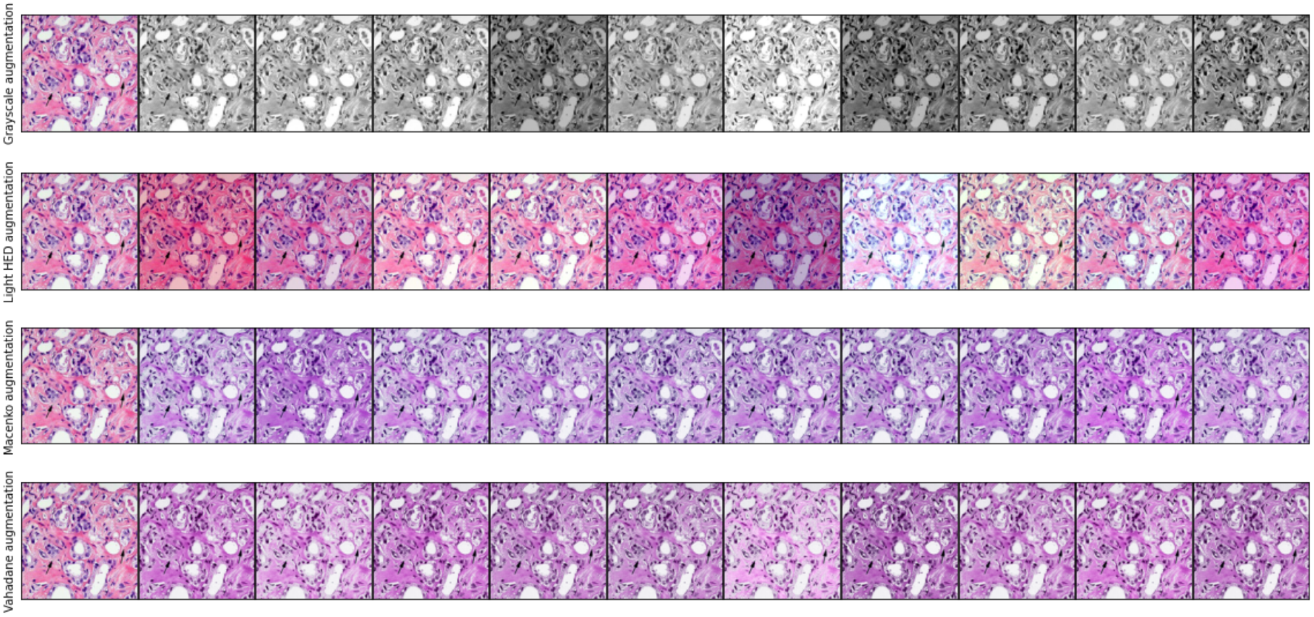
Illustrative examples for the augmentation methods of stainlib

### 3.1. Image Normalization

Figure 5 and 6 show the results for the Vahadane and Macenko normalizations methods. Both methods estimate the stain concentration matrix, Vahadane with a non-negative matrix factorization approach and Macenko with a singular value decomposition. Thus, results appear similar with the images normalized with Macenko being slightly darker or with more stain concentration than with the Vahadane method. In Figure 7, examples for the Reinhard normalization method are presented. Because the method aims to match the color histogram of the target image directly, the background matches the lightest color in the target image, which produces unrealistically looking images. We included a tissue detector that masks the tissue content from the background to alleviate this. The tissue detector is included as a parameter in the call for transforming new images using the Reinhard method in `stainlib` ^3^.

**Figure 5:**
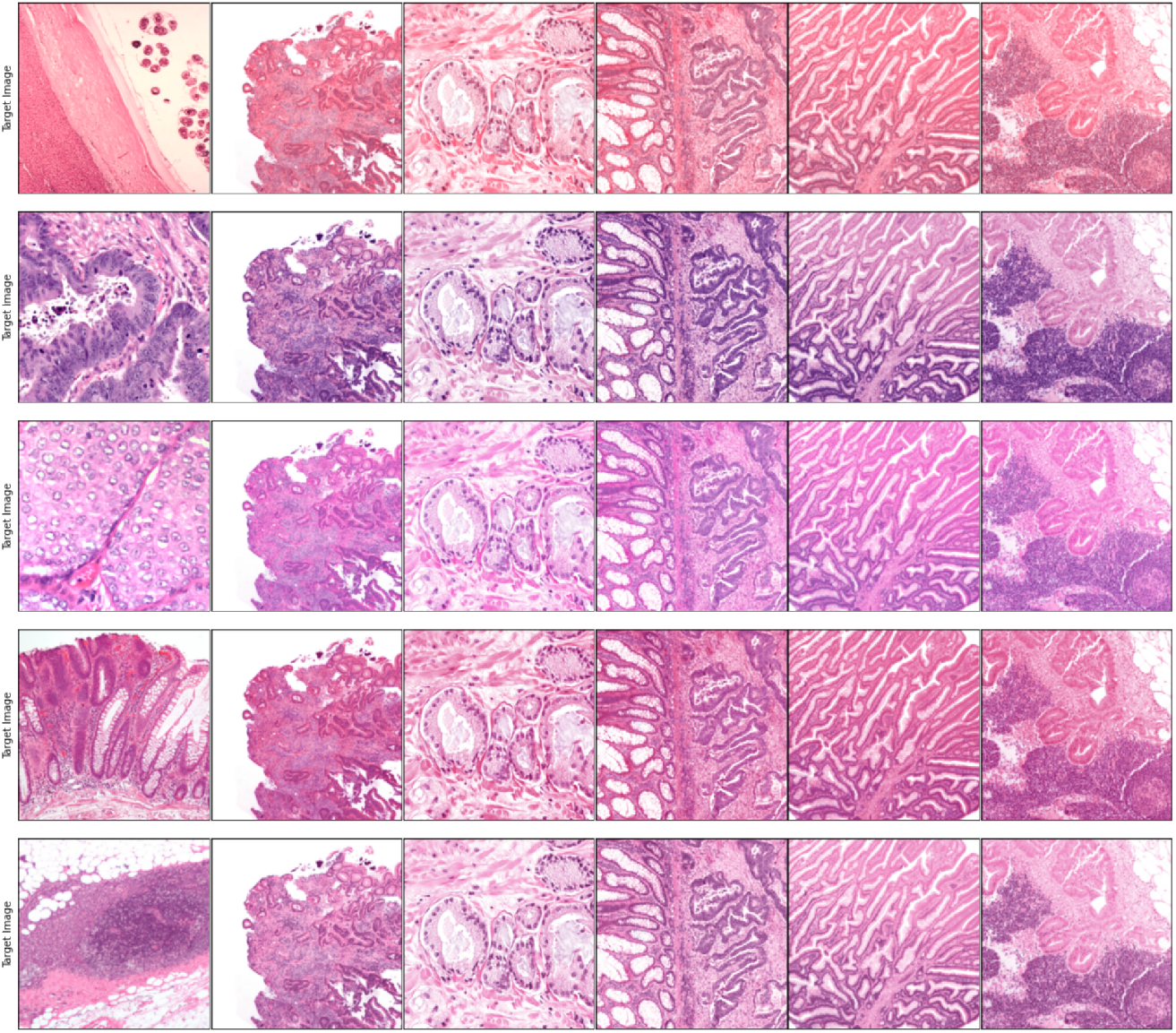
Illustrative examples for the implemented Vahadane normalization method in stainlib.

**Figure 6:**
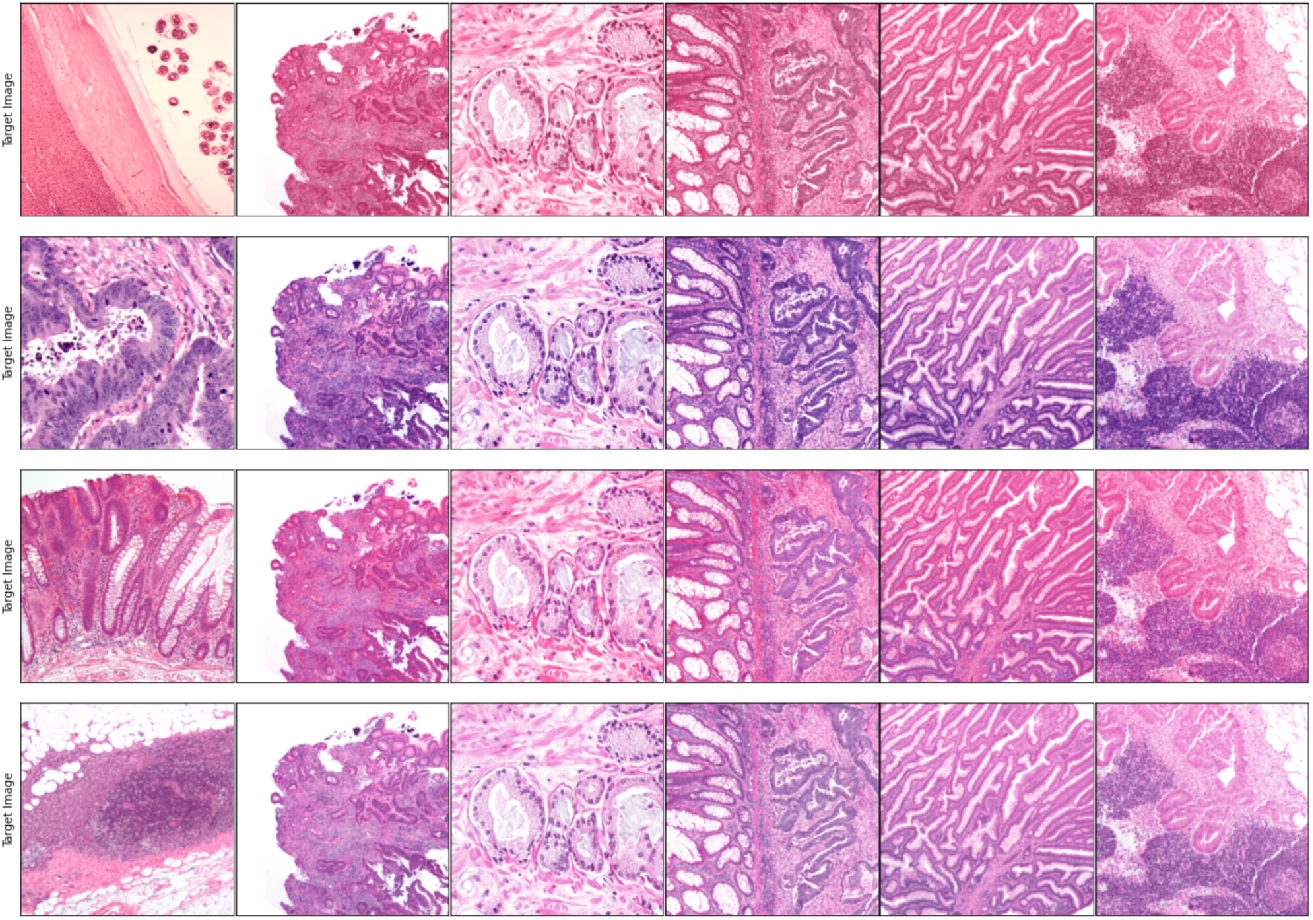
Illustrative examples for the implemented Macenko normalization method in stainlib.

**Figure 7:**
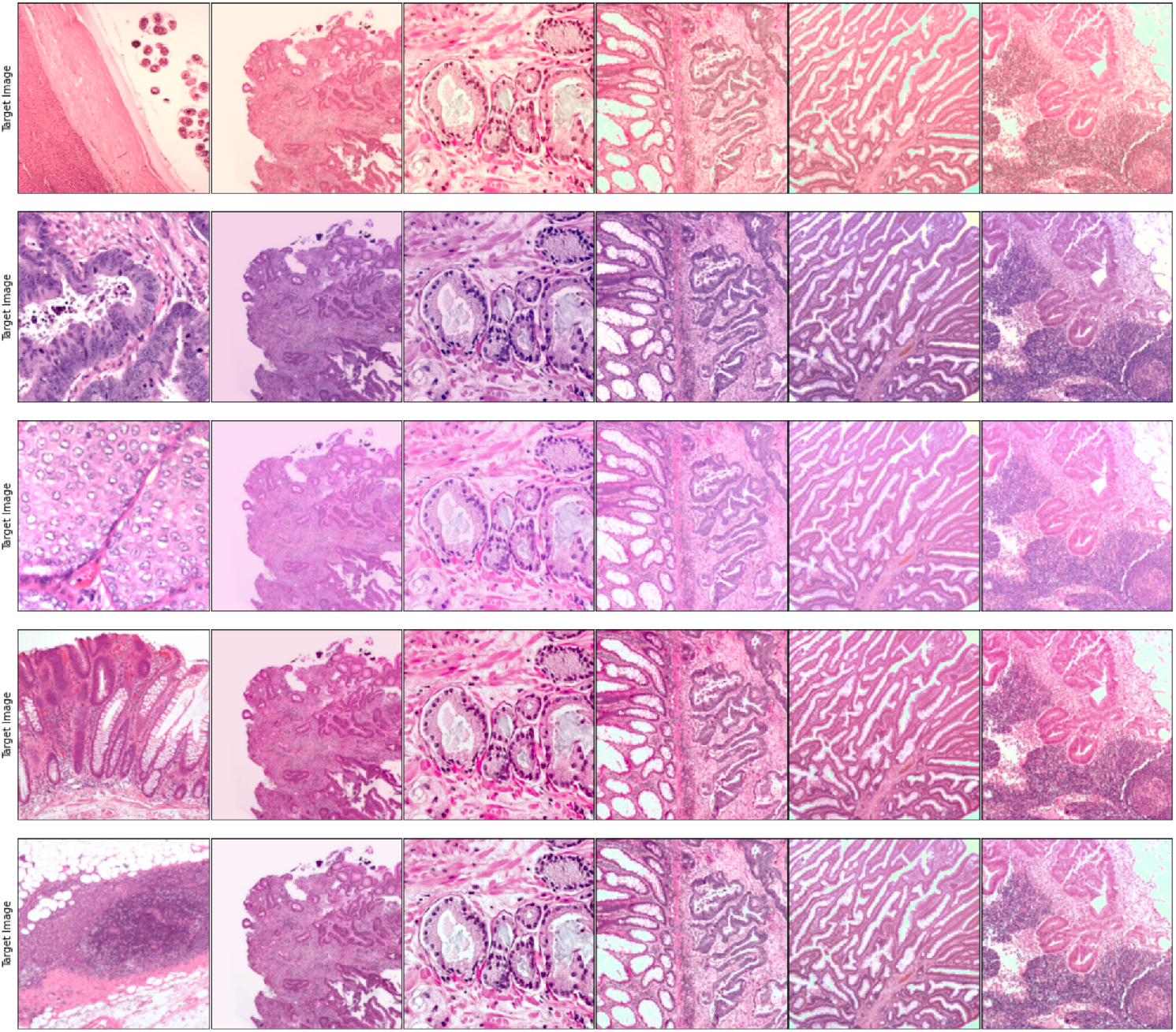
Illustrative examples for the implemented Reinhard normalization method in stainlib.

## 4. Impact

`stainlib` allows researchers worldwide to tackle scientific challenges in computational pathology more easily in several ways. First, `stainlib` allows testing new algorithms under multiple data-heterogeneity scenarios, given the characteristics of the augmented and normalized images that simulate uncontrolled variations in the real-world test sets. For example, it allows researchers to test an algorithm having specific performance on a given test set on realistic variations of the same dataset generated with `stainlib`, to test if the performance remains the same over the variations or the algorithms requires additional improvement (potentially including such variations of the dataset in the training data). Second, it empowers researchers to use small data sets for training deep learning models by augmenting significantly the data sets with color-shifted versions of the training data. The bigger data-augmented training sets make trained CNN models more robust and less prone to errors. Third, `stainlib` includes very recent image processing and machine learning techniques reported in the literature to deal with color heterogeneity. Finally, the `stainlib`’s modular structure is designed to be easy to expand and the authors encourage researchers to contribute to this library by suggesting changes and improvements or directly adding or improving methods into the code base.

## 5. Conclusions

This article presents and qualitatively tests the `stainlib` library, implemented to include novel and widely used tools that extract homogeneous representations of heterogeneous color visual information from H&E images. `stainlib` is easy to use and to expand and it includes the most commonly used methods not only for stain normalization but also for augmentation, together with recent deep learning based approaches. We anticipate continuous updates for `stainlib`, making it efficient and useful for normalizing image regions and WSIs. We also strongly encourage the contribution of additional tools by researchers worldwide. The source code for all the tools is now fully accessible in the repositories. We are confident that these resources allow researchers to build more easily and quickly robust models that generalize to unseen images from heterogeneous sources.

**Table 1:**
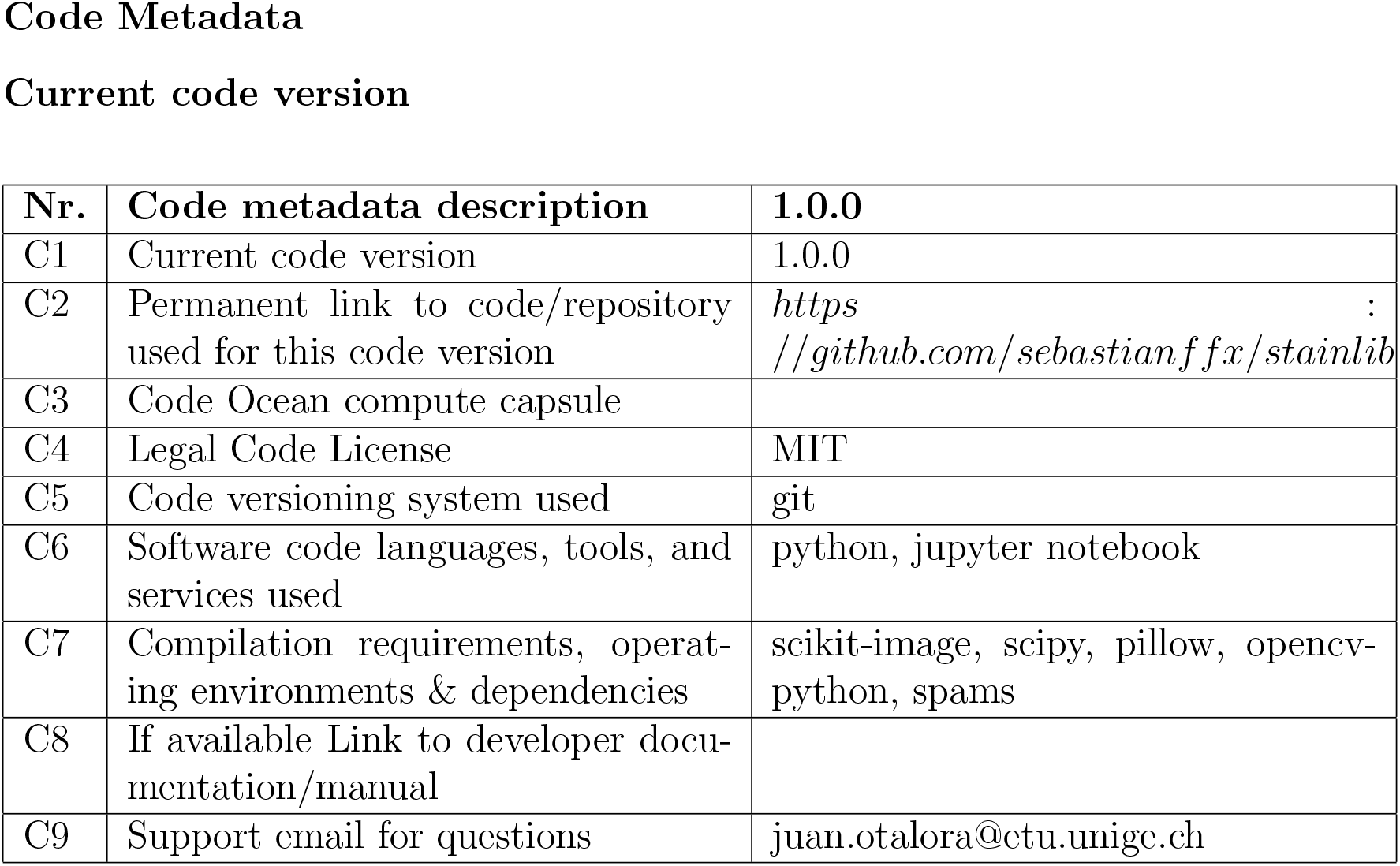
Code metadata (mandatory).

## Acknowledgements

We developed `stainlib` as part of the ExaMode project^4^. This project has received funding from the European Union’s Horizon 2020 research and innovation programme under grant agreement No 825292 (ExaMode, http://www.examode.eu/). Sebastian Otálora thanks Minciencias through the call 756 for PhD studies.

## Current executable software version

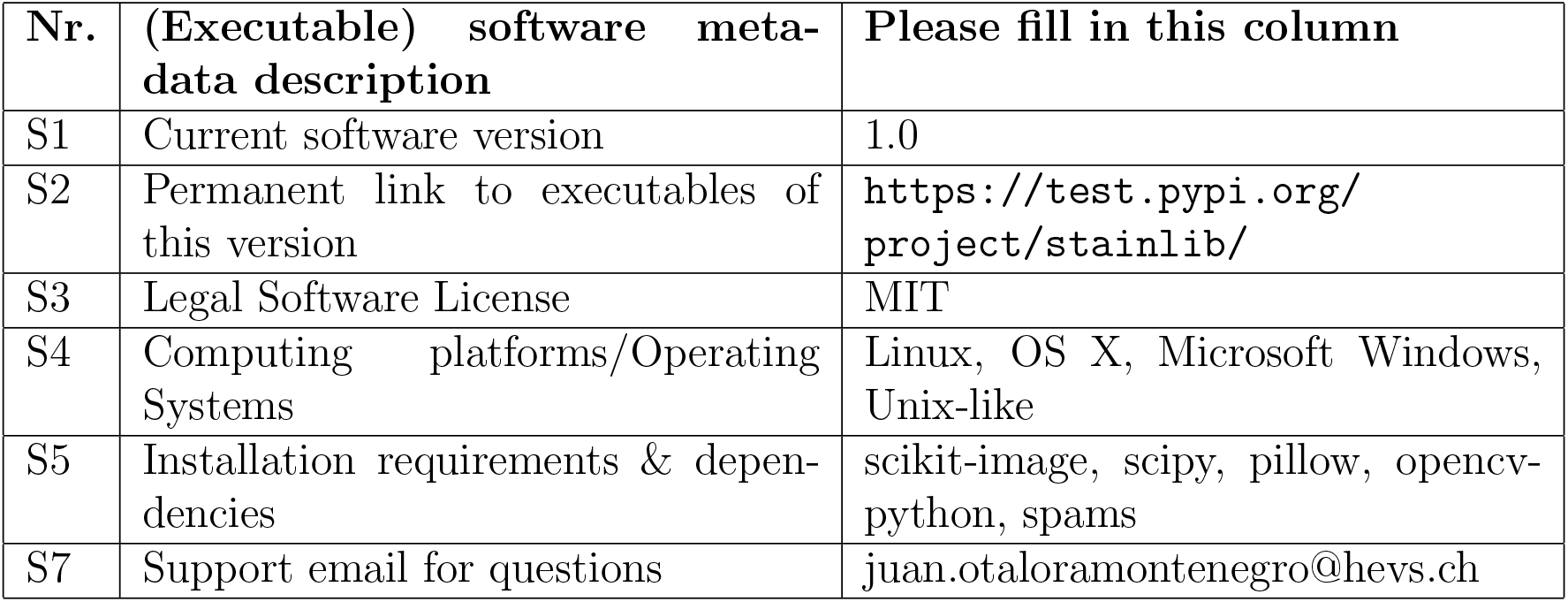

## Footnotes

1 Staintools github repository: https://github.com/Peter554/StainTools/tree/master/staintools

2 2Invertible ows github repository:https://github.com/sara-nl/color-information

3 https://github.com/sebastianffx/stainlib/blob/main/normalization/normalizer.py

4 https://www.examode.eu/

